# Quantitative proteomic analysis of auxin signaling during seedling development

**DOI:** 10.1101/211532

**Authors:** Dior R. Kelley, Zhouxin Shen, Justin W. Walley, Elisabeth J. Chapman, Steven P. Briggs, Mark Estelle

## Abstract

Auxin induces rapid gene expression changes throughout plant development. How these transcriptional responses relate to changes in protein abundance is not well characterized. We have identified auxin regulated proteins in whole seedlings, roots and hypocotyls and at three different time points (30 min, 120 min and 3 hours) using an iTRAQ (isobaric tags for relative and absolute quantification) based quantitative proteomics approach. These profiling experiments detected 4,701 proteins from seedling tissue, 6,740 proteins from root tissue and 3,925 proteins from hypocotyl tissue. Comparisons between the differentially expressed proteins data sets showed little overlap, suggesting that auxin proteomes exhibit both temporal and spatial specificity. Numerous proteins showed significant changes in abundance following auxin treatment independent of changes in cognate transcript abundance. This includes several well characterized proteins with various roles in auxin pathways, suggesting that complex gene regulation mechanisms follow auxin signaling events. Specifically, regulation of translation may play a role as inferred from MapMan categorization analyses and protein interaction networks comprised of auxin regulated proteins. Additionally, functional categorization of auxin regulated proteins indicates rapid and complex metabolic changes occur in both roots and hypocotyls in response to auxin which are not apparent from transcriptome analyses. Altogether these data describe novel auxin-regulated proteomes and are an excellent resource for identifying new downstream signaling components related to auxin-mediated plant growth and development.

## Introduction

Auxin is one of the major phytohormones involved in regulating many aspects of seedling development, including cotyledon formation, hypocotyl cell elongation, meristem maintenance and root morphogenesis (Finet and Jaillais, 2012). In land plants, the response to auxin is controlled by co-receptors comprised of TIR1/AUXIN F-BOX (TIR1/AFB) and Aux/IAA transcriptional regulators. In *Arabidopsis* there are 6 TIR1/AFB proteins and 29 Aux/IAA proteins (Strader and Zhao, 2016). The Aux/IAA proteins actively repress transcription by interacting with transcription factors called AUXIN RESPONSE FACTORS (ARFs) transcription factors and recruiting a co-repressor protein, TOPLESS (TPL). Auxin acts by promoting the degradation of the Aux/IAAs leading to tightly regulated gene expression changes that have been well documented (Bargmann et al., 2014; Chapman et al., 2012; Overvoorde et al., 2005; Weijers and Wagner, 2016).

One of the outstanding questions in the field is how these auxin-mediated transcriptional changes collectively influence proteome composition. Arabidopsis seedlings are an excellent model for comparative proteomics because they exhibit tissue-specific auxin responses and provide sufficient quantities of plant material for sampling. Additionally, transcriptional changes in Arabidopsis seedlings, hypocotyls and roots have been well documented (Bargmann et al., 2014; Chapman et al., 2012; Laskowski et al., 2006; Lewis et al., 2013; Nemhauser et al., 2006; Overvoorde et al., 2005; Stepanova et al., 2007). Initial characterization of auxin responsive proteomes in seedlings and roots identified proteins that are responsive 6-24 hours after auxin treatment (Slade et al., 2017; Xing and Xue, 2012) and protein phosphorylation events associated with auxin mediated lateral root formation (Zhang et al., 2013). Further studies of the auxin-regulated proteome are needed to generate a more comprehensive view of auxin mediated gene expression (Mattei et al., 2013).

In this study, we characterized the auxin-regulated proteome in Arabidopsis seedlings after 3 hrs of auxin treatment as well as in dissected hypocotyls and roots 30 and 120 minutes after exposure to auxin. These data sets provide a proteomic description of how auxin influences gene expression in a temporal and spatial fashion. Comparisons between these proteomes showed little-to-no overlap in the differentially expressed proteins, suggesting that organ-specific auxin responses are mediated by distinct cellular proteomes. Additionally, these data can be used to assess the relevance of auxin mediated transcriptional responses and potentially uncover novel proteins downstream of co-receptor action that may mediate diverse developmental programs.

## Material & Methods

### Plant Material

*Arabidopsis thaliana* plants used in this study were *Columbia* (Col-0) ecotype. Seeds were surfaced sterilized using 50% bleach and 0.01% Triton X-100 for 10 minutes and then washed five times with sterile water. Seeds were then imbibed in sterile water for two days at 4°C and then transferred to 0.5X Murashige-Skoog medium plates overlaid with sterile nylon mesh squares. Seedlings were grown under long day photoperiods (16 h light/8 h dark) at 23°C. Auxin treatments on hypocotyls were done as previously described (Chapman et al., 2012). Five day-old seedlings were treated with DMSO (mock), IAA or picloram (auxin) as specified. For seedling profiling, intact seedlings were treated and immediately frozen in liquid nitrogen; seedlings were pooled to reach 1 g of tissue per biological replicate. For root profiling, intact seedlings were treated and roots were carefully dissected from the rest seedling using a scalpel then immediately frozen; roots were pooled to reach 1 g of tissue per biological replicate. For hypocotyl profiling, intact seedlings were treated and hypocotyls were carefully dissected from the rest seedling using a scalpel then immediately frozen into liquid nitrogen; hypocotyls were pooled to reach 0.5 g tissue per biological replicate. Four independent biological replicates were generated for seedling and root treatments while two replicates were collected for hypocotyls due to the limited nature of this tissue.

### Preparation and Analysis of Proteins via Mass Spectrometry

Peptide preparation, and protein abundance profiling by mass spectrometry are based on previously described methods (Walley et al., 2013, 2015). Each frozen tissue sample was thoroughly ground to a fine powder for 15 min in liquid nitrogen prior to protein extraction. Proteins were precipitated and washed with 50 ml of -20 °C methanol three times, then 50ml -20 °C acetone three times. Protein pellets were aliquoted into four 2 ml Eppendorf tubes and dried in a vacuum concentrator at 4 °C. Protein pellets were suspended in 1 ml extraction buffer (0.1% SDS, 1 mM EDTA, 50 mM Hepes buffer, pH 7). Cysteines were reduced and alkylated using 1 mM Tris (2-carboxyethyl)phosphine (Fisher, AC36383) at 95 °C for 5 minutes, then 2.5 mM iodoacetamide (Fisher, AC12227) at 37°C in the dark for 15 minutes, respectively. Protein was quantified using a Bradford assay with bovine serum albumn used to construct the standard curve (Pierce). Proteins were digested with trypsin overnight (Roche, 03 708 969 001, enzyme:substrate w:w ratio = 1:100). A second digestion was performed the next day for 4 hours (enzyme:substrate w:w ratio = 1:100). Digested peptides were purified on a 500 mg Waters Oasis MCX cartridge to remove SDS. Peptides were eluted from the MCX column with 4 ml 50% isopropyl alcohol and 400 mM NH_4_HCO_3_ (pH 9.5) and then dried in a vacuum concentrator at 4 °C. Peptides were resuspended in 0.1% formic acid and further purified on a 50 mg Sep-Pak C18 column (Waters). Peptide amount was quantified using the Pierce BCA Protein assay kit with bovine serum albumn used to construct the standard curve.

Peptides were labeled with iTRAQ reagents (AB SCIEX) and we obtained higher than 95% iTRAQ labeling efficiency by treating 100 µg of non-modified peptides with one tube of iTRAQ reagent for 2 hours at room temperature. Labeled samples were dried down in a vacuum concentrator and resuspended in 0.1% formic acid. Samples tagged with the four different iTRAQ reagents were pooled together.

An Agilent 1200 HPLC system (Agilent Technologies) was used to deliver a flow rate of 600 nL min^-1^ via a 3-phase capillary chromatography column through a splitter to the mass spectrometer. The 3-phase capillary chromatography was set up as follows. Using a custom pressure cell, 5 µm Zorbax SB-C18 (Agilent) was packed into fused silica capillary tubing (200 µm ID, 360 µm OD, 30 cm long) to form the first dimension reverse-phase column (RP1). A 5 cm long strong cation exchange (SCX) column packed with 5 µm PolySulfoethyl (PolyLC) was connected to RP1 using a zero dead volume 1 µm filter (Upchurch, M548) attached to the exit of the RP1 column. A fused silica capillary (200 µm ID, 360 µm OD, 20 cm long) packed with 2.5 µM C18 (Waters) was connected to SCX as the analytical column (RP2). The electrospray tip of the fused silica tubing was pulled to a sharp tip with the inner diameter smaller than 1 µm using a laser puller (Sutter P-2000). The peptide mixtures were loaded onto the RP1 column using the custom pressure cell. A new set of columns was used for each LC-MS/MS analysis.

Peptides were first eluted from the RP1 column to the SCX column using a 0 to 80% acetonitrile gradient for 150 minutes. The peptides were then fractionated by the SCX column using a series of 28 salt steps for non-modified iTRAQ profiling (0, 20, 40, 50 55, 60, 62.5, 65, 67.5, 70, 72.5, 75, 77.5, 80, 82.5, 85, 87.5, 90, 92.5, 95, 97.5, 100, 120, 150, 180, 200, 500, 1000 mM ammonium acetate) followed by high-resolution reverse phase separation using an acetonitrile gradient of 0 to 80% for 120 minutes.

Spectra were acquired using an LTQ Velos linear ion trap tandem mass spectrometer (Thermo Electron Corporation, San Jose, CA) employing automated, data-dependent acquisition. The mass spectrometer was operated in positive ion mode with a source temperature of 250 °C. The full MS scan range of 400-2000 m/z was divided into 3 smaller scan ranges (400-800, 800-1050, 1050-2000) to improve the dynamic range. Both CID (Collision Induced Dissociation) and PQD (Pulsed-Q Dissociation) scans of the same parent ion were collected for protein identification and quantitation. Each MS scan was followed by 4 pairs of CID-PQD MS/MS scans of the most intense ions from the parent MS scan. A dynamic exclusion of 1 minute was used to improve the duty cycle of MS/MS scans.

The raw data were extracted and searched using Spectrum Mill v3.03 (Agilent). The CID and PQD scans from the same parent ion were merged together. MS/MS spectra with a sequence tag length of 1 or less were considered to be poor spectra and were discarded. The remaining MS/MS spectra were searched against the *Arabidopsis* TAIR10 database. The enzyme parameter was limited to fully tryptic peptides with a maximum miscleavage of 1. All other search parameters were set to default settings of Spectrum Mill (carbamidomethylation of cysteines and iTRAQ modification). A concatenated forward-reverse database was constructed to calculate the in-situ false discovery rate (FDR). All datasets were summarized together to maintain FDR across the datasets. Cutoff scores were dynamically assigned resulting in FDR of 0.05%, 0.12%, and 0.66% at the spectrum, peptide, and protein level, respectively. Proteins that share common peptides were grouped to address the database redundancy issue. The proteins within the same group shared the same set or subset of unique peptides.

iTRAQ intensities were calculated by summing the peptide iTRAQ intensities from each protein group. Peptides shared among different protein groups were removed before quantitation using custom Perl scripts implemented in Spectrum Mill v3.03 (Agilent). Isotope impurities of iTRAQ reagents were corrected using correction factors provided by the manufacturer (Applied Biosystems). Median normalization was performed to normalize the protein iTRAQ reporter intensities in which the log ratios between different iTRAQ tags (115/114, 116/114, 117/114) are adjusted globally such that the median log ratio is zero. Protein ratios between the mock and each treatment were calculated by taking the ratios of the total iTRAQ intensities from the corresponding iTRAQ reporter.

Protein ratios were then log_2_ converted. Proteins that significantly changed in each treatment, relative to mock, were determined using t-tests (two-sample heteroscedastic). Proteins with a p-value of <0.05 were considered to be significantly differentially expressed.

### GO enrichment and MapMan analysis

GO enrichment analyses were performed using Gorilla (<u>http://cbl-gorilla.cs.technion.ac.il/)</u> (Eden et al., 2009). Functional categorization of differentially expressed proteins was performed using the Classification SuperViewer Tool with the classification source set to MapMan (<u>http://bar.utoronto.ca/)</u> (Provart et al., 2003).

### STRING network analyses

Protein-protein interaction networks were generated using STRING (<u>https://string-db.org/)</u> (Szklarczyk et al., 2017). Differentially expressed proteins were used as a multiple search by protein names/identifiers against Arabidopsis thaliana as the organism.

## Results

### Quantitative proteomic analysis of seedlings following auxin treatments indicate tissues specific responses

The transcriptional response to exogenous auxin in various tissues and species have been well characterized. We sought to define rapid proteome changes in young seedlings, roots and hypocotyls using peptide mass spectrometry. The organs and time points selected have been well characterized transcriptionally and are developmentally relevant. Following auxin treatment, light-grown hypocotyls elongate while primary root elongation is inhibited. Thus, we hypothesized that we would uncover protein abundance changes specific to each organ profiled. Towards this aim we treated seedlings with a naturally occurring auxin, indole-3-acetic acid (IAA) and harvested both whole seedlings and intact roots for proteome profiling. Additionally, we treated seedlings with a synthetic auxin, picloram, and then harvested hypocotyls for proteome profiling. Two different auxins were used in our workflow because we wanted to be able to directly compare the corresponding proteome datasets to previously published transcriptomic data generated in the same fashion (Chapman et al., 2012; Lewis et al., 2013; Nemhauser et al., 2006). Following auxin and mock solvent control treatments, total protein was extracted from these samples and digested into tryptic peptides. Peptides were iTRAQ labeled and then identified and quantified using tandem mass spectrometry (MS/MS) (Figure 1A). For the root and seedling samples, four biological replicates were generated for each condition. Because it is difficult to collect large amounts of Arabidopsis hypocotyl tissue, only two biological replicates were generated for each condition.

**Figure 1.**
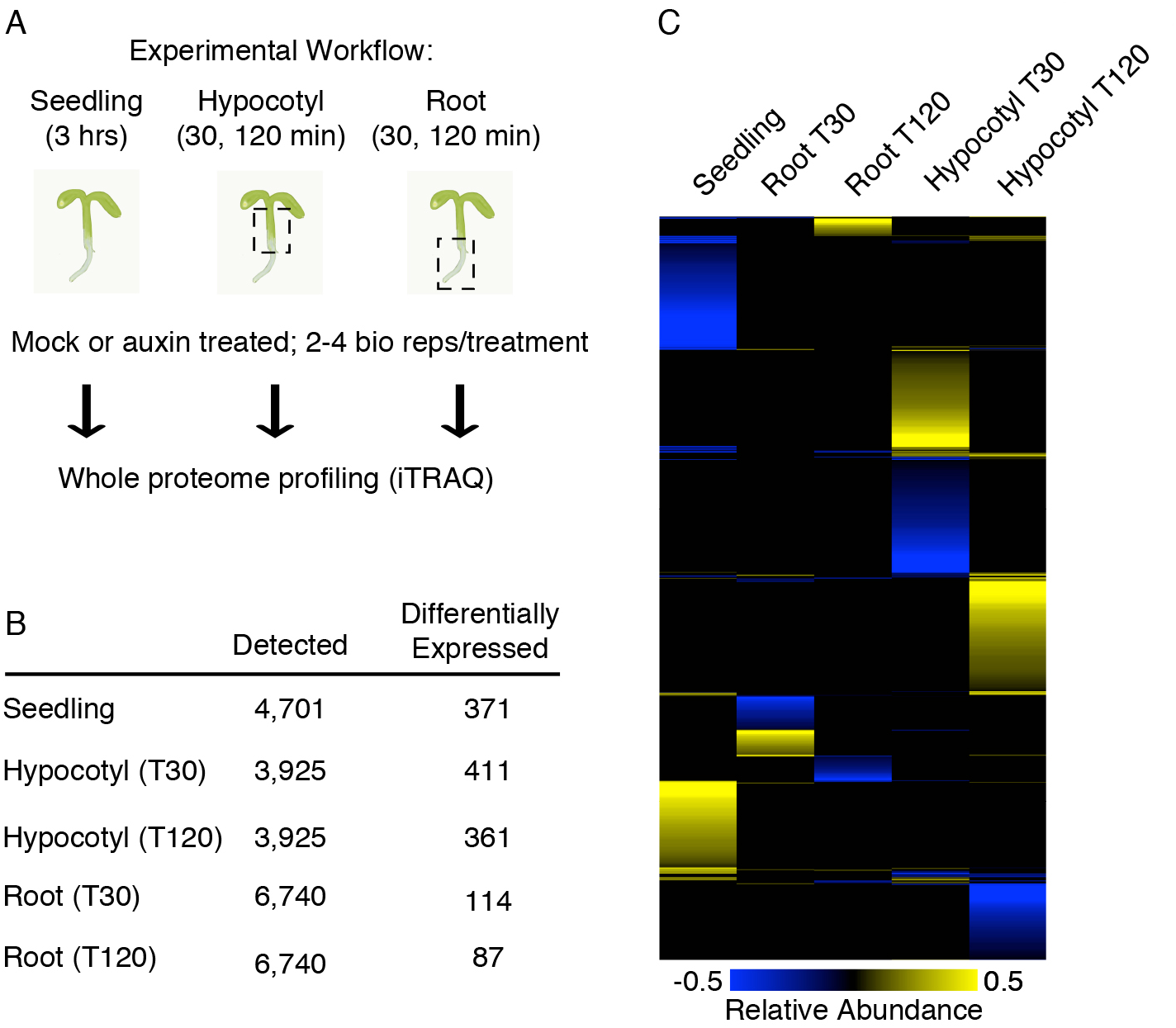
Quantitative proteomic analysis of seedlings following auxin treatment demonstrates tissues specific responses. (A) Schematic of the experimental workflow. Five day-old wild-type seedlings were treated with auxin or a mock solution for the specified time period. Intact seedlings were harvested after 3 hours of treatment. Hypocotyls and roots were dissected away from treated seedlings either 30 minutes or 120 minutes after treatment. Whole proteome profiling was performed on the samples using iTRAQ. (B) The total number of proteins detected in each tissue sampled and the number exhibiting differential expression in the auxin treated samples relative to mock treated samples, p-value <0.05. (C) Hierarchical clustering of differentially expressed proteins shows several distinct clusters between tissue types and time points.

A total of 4,701 proteins were detected in the seedling samples of which 371 changed in abundance after auxin treatment (*P* ≤ 0.05) (Figure 1B; Table S1). From the hypocotyl samples 3,925 proteins were detected while 411 proteins were significantly changed at 30 min and 361 proteins were significantly changed at 120 min (Figure 1B; Table S1). From the root samples 6,740 proteins were detected while 114 proteins were significantly changed at 30 min while 87 proteins were differentially expressed at 120 min (Figure 1B; Table S1). In order to examine the extent of tissue specificity, we performed hierarchical clustering with all of the differentially expressed proteins from all tissues and time points sampled (*P*-value <0.05). Each set of differentially expressed proteins from the different tissues that were sampled is fairly discrete and we observed little overlap among these datasets (Figure 1C). Very few differentially regulated proteins appear to be in common between the seedling, hypocotyl and root proteomes. This may be due to our limited sampling of the genome, as we did not detect more than ~6,700 proteins from any run (Figure 1B; Table S1). It could also indicate that the observed auxin proteomes have a rapid and transient nature. However, there were some proteins that were differentially expressed within an organ at both time points sampled and some proteins in common between organs at the same time point. For example, within the hypocotyl there are 14 differentially expressed proteins in common to both time points (Figure 2A). Such proteins include RUB1 CONJUGATING ENZYME (RCE1; At4g36800) which has known roles in auxin mediated processes (Dharmasiri et al., 2003; Esteve-Bruna et al., 2013; Pozo, 1998) and SNF7.1 (At4g29160) a member of the ESCRT-III complex involved in vesicle formation (Cai et al., 2014). Altogether the observed protein abundance profiles following auxin exposure exhibit tissue specific responses.

**Figure 2.**
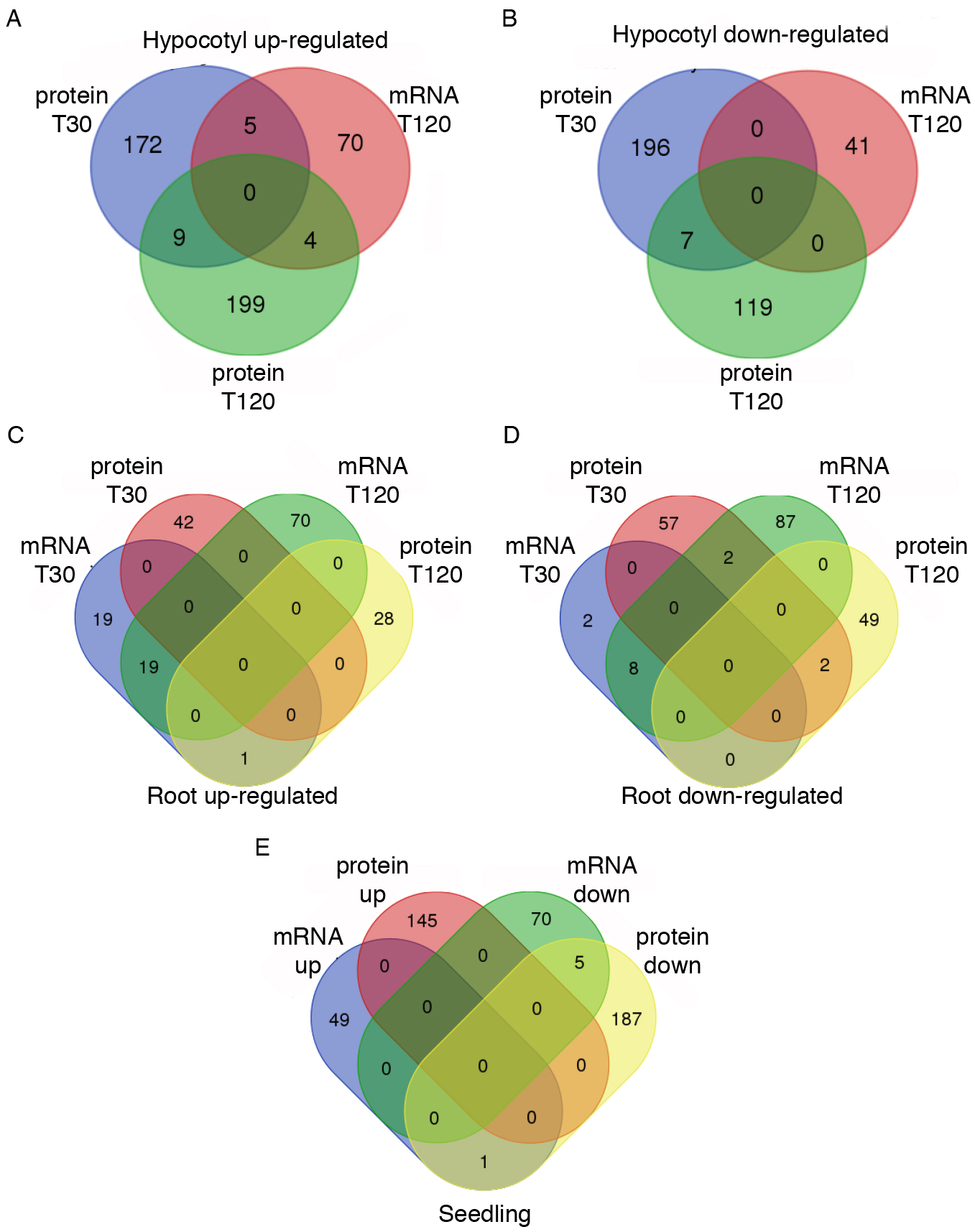
Auxin regulated transcriptomes are discordant with auxin regulated proteomes. (A) Venn diagram comparing all of the hypocotyl up-regulated proteins with published transcriptome data. (B) Venn diagram comparing all of the hypocotyl down-regulated proteins with published transcriptome data. (C) Venn diagram comparing all of the root up-regulated proteins with published transcriptome data. (D) Venn diagram comparing all of the root down-regulated proteins with published transcriptome data. (E) Venn diagram comparing all of the up and down auxin regulated proteins in seedlings with published transcriptome data.

### Auxin regulated transcriptomes do not correspond with proteomes

A number of studies have shown weakly positive correlation between transcriptome and proteome changes among eukaryotic organisms (Baerenfaller et al., 2008; Ghaemmaghami et al., 2003; Ghazalpour et al., 2011; Ponnala et al., 2014; Schwanhausser et al., 2011; Vogel et al., 2010; Walley et al., 2013, 2016; Washburn et al., 2003). In order to examine the extent of correspondence between auxin regulated transcriptional responses and the proteome changes observed in this study we compiled a list of genes for which both mRNA and protein were detected (Chapman et al., 2012; Lewis et al., 2013; Nemhauser et al., 2006). From the seedling datasets (Nemhauser et al., 2006) 4,343 genes were detected at both the mRNA and protein levels. From the hypocotyl datasets (Chapman et al., 2012) 3,888 genes were detected at both the mRNA and protein levels. From the root datasets (Lewis et al., 2013) 4,929 genes were detected at both the mRNA and protein levels.

Specifically, only five genes are decreased in abundance at both the mRNA and protein level in seedlings following auxin treatment (Figure 2E). This includes a member of the CYP705A family of cytochrome P450 enzymes (AT5G47990) that plays a role in gravitropism (Withers et al., 2013). Notably, one gene has increased mRNA but decreased protein abundance (AT5G54500). AT5G54500 encodes a flavodoxin-like quinone reductase that is annotated as a primary auxin-response gene. One explanation for this observed response may be that the AT5G54500 protein is not stable and is rapidly degraded following auxin treatment. In the hypocotyl, only five genes are upregulated at both the mRNA and protein level while the downregulated genes and proteins are discordant (Figure 2A,B). A similar pattern was observed among the root regulated genes and proteins (Figure 2 C,D). Overall, these comparisons demonstrate that there is very little overlap between measured auxin mediated transcriptional responses and proteome responses (Figure 2).

### Key auxin responsive proteins are observed across tissue types

We examined the observed differentially expressed proteins in more detail in order to identify particular proteins that may play known roles in auxin biology. Numerous proteins with altered abundance levels in auxin-treated seedlings have been previously linked to auxin pathways, providing support for these profiling data (Figure 3 and Table S1). Such proteins include the auxin-responsive GH3 family protein GH3.17 (Gonzalez-Lamothe et al., 2012; Staswick, 2005), the auxin influx carrier LIKE AUXIN RESISTANT 1 (LAX1) (Swarup et al., 2004; Ugartechea-Chirino et al., 2010), auxin efflux carrier PIN-FORMED 3 (PIN3) (Friml et al., 2002; Paponov et al., 2005), AUXIN RESPONSE FACTOR 7 (ARF7)/MASSUGU1 (MSG1) (Harper et al., 2000; Watahiki et al., 1999), and ARF8 (Tian et al., 2004), PROTEIN PHOSPHATASE 2A A3 (PP2AA3) which regulates auxin efflux (Dai et al., 2012) and (RELATED TO UBIQUITIN) RUB1 which plays a key role in modulating SCF assembly and thus auxin receptor activity by altering AtCUL1 modification (del Pozo et al., 2002; Pozo, 1998). Notably, a number of these proteins do not have corresponding transcripts that are auxin responsive.

**Figure 3.**
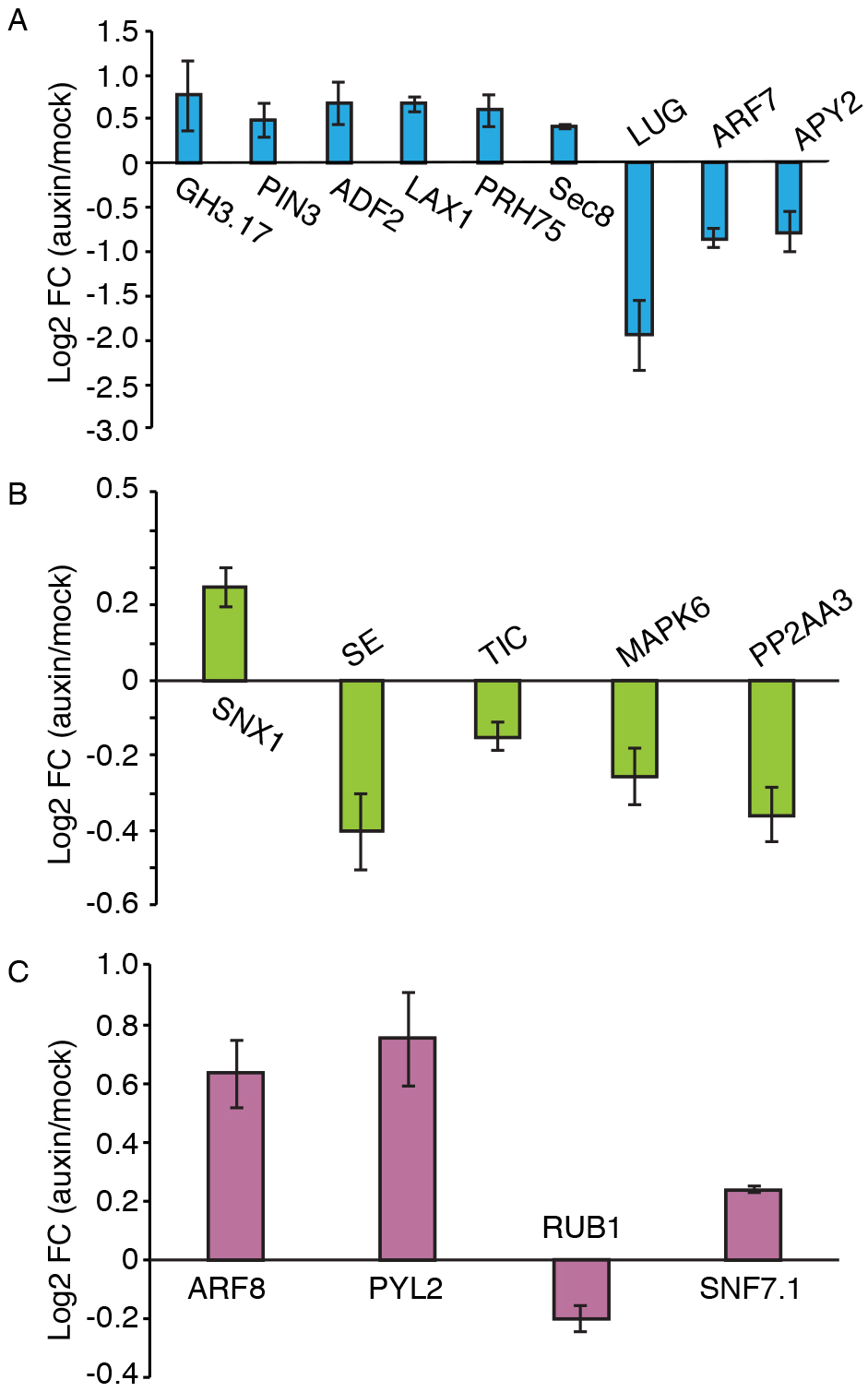
Key proteins known to be involved in auxin pathways exhibit discrete protein abundance profiles. (A) In the seedling, proteins such as GH3.17, PIN3, ADF2, LAX1, PRH75 and Sec8 are upregulated while LUG, ARF7 and APY2 are downregulated in response to auxin. (B) In the root, SNX1 is upregulated while SE, TIC, MAPK6 and PP2AA3 are downregulated following auxin treatment. (C) In the hypocotyl, ARF8,PYL2 and SNF7.1 are upregulated while RUB1 is decreased in abundance following picloram (auxin) treatment.

### Biological process and function ontologies varied between auxin-regulated proteomes

We hypothesized that the functional classification of proteins involved in auxin mediated responses may vary between organs. In order to explore this idea further we performed GO enrichment analyses by comparing the differentially expressed proteins against all the proteins detected (Table S1). These results show different parent terms overrepresented among the different tissues and time points sampled but there were no striking enrichment terms (Table S2). Using a different approach, we put each list of differentially expressed proteins into the classification SuperViewer tool from BAR (Waese and Provart, 2016) in order to examine the categorical distribution of differentially expressed proteins within each dataset (Figure 4). For each dataset examined, we observed several functional classifications with normalized class scores >1, consistent with the notion that auxin signaling can affect many downstream cellular processes. Examples of common categories include redox, amino acid metabolism, stress (both biotic and abiotic), protein, RNA processing, hormone metabolism, lipid metabolism, transport, signaling and secondary metabolism. The “proteins” functional category is predominantly comprised of proteins associated with translation and synthesis.

**Figure 4.**
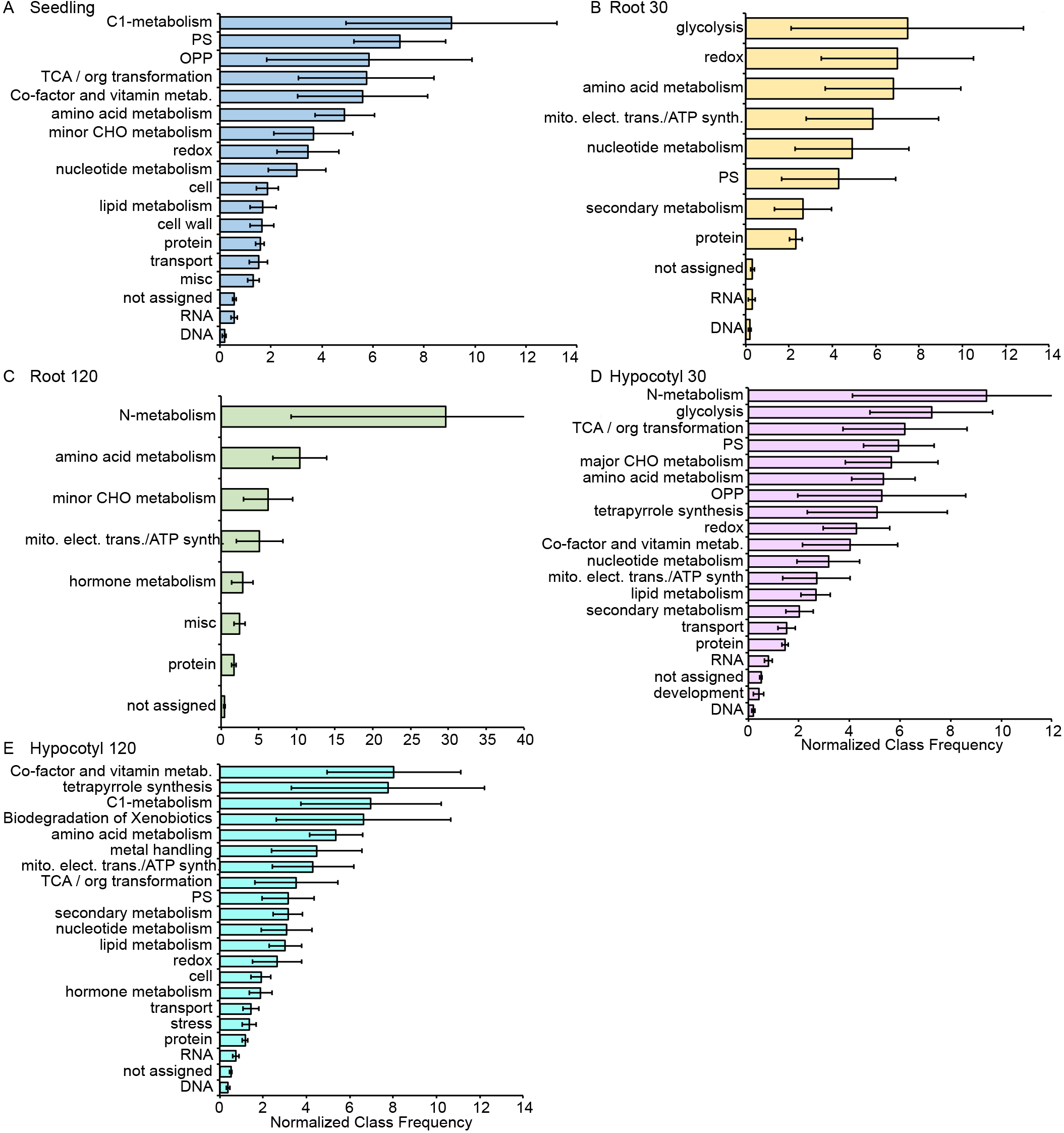
Auxin regulated proteins can be classified into diverse functional MapMan categories and exhibit a common categorical signature. (A) Upon auxin treatment differentially expressed proteins detected in seedlings (A), roots (B,C) and hypocotyls (D,E) fall into several common functional categories including amino acid metabolism, redox and nitrogen metabolism.

Notably hormone metabolism is affected differently between organs and time points, with auxin being common to most organs and time points and brassinosteroid (BR), abscisic acid (ABA), jasmonic acid (JA) and ethylene co-occurring in many instances (Figure 4). However, ethylene metabolism appears to specifically be downregulated in both hypocotyls and roots. JA metabolism is differentially regulated in hypocotyls but not roots. ABA metabolism was not affected in seedlings but was differentially regulated in hypocotyls and roots.

A few other categories exhibit striking patterns across these data sets. Proteins associated with mitochondrial electron transport are solely associated with downregulated proteins in all the organs/time points. Proteins involved with cell vesicle transport appear to be upregulated in hypocotyls at 30 min and roots at 120 min. Proteins falling into the “development” functional category are only observed as changing in seedlings.

### Protein interaction networks underlying auxin regulated root proteomes

We examined protein interaction networks generated from the tissue-specific datasets in order to uncover higher level connections and identify processes that may be relevant to auxin signaling (Figure 5 and data not shown). In order to examine interactions among proteins exhibiting altered abundance in response to auxin we performed STRING analysis (Franceschini et al., 2013) on the differentially expressed seedling, hypocotyl and root datasets. STRING networks are generated from known and predicted protein-protein interactions, which can include direct physical interactions and indirect functional associations (Franceschini et al., 2013; Szklarczyk et al., 2015). Shown in Figure 5 are the STRING results from the auxin regulated differentially expressed proteins detected in roots at both time points. Both datasets displayed a high degree of connectivity, only the root protein-protein interaction networks are shown in Figure 5. This high degree of connectivity suggests that the differentially expressed proteins identified may be biologically connected as a group (Figure 5 and data not shown). At both time points the root auxin proteomes consisting of 114 proteins (T30) and 87 proteins (T120) had significantly more interactions than expected; 251 edges observed compared to 62 expected and 93 edges observed compared to 34 expected, for T30 and T120, respectively (Figure 5). Notably, the central clusters observed in both root networks were dominated by ribosomal subunits and elongation factors; “translation” GO enrichment value FDR = 8.79e-09 (T30) and 0.0439 (T120). This is consistent with transcriptional and mutant studies indicating that various auxin signaling pathways are under translational control (Horiguchi et al., 2012; Nishimura et al., 2005; Rosado et al., 2012; Zhou et al., 2010).

**Figure 5.**
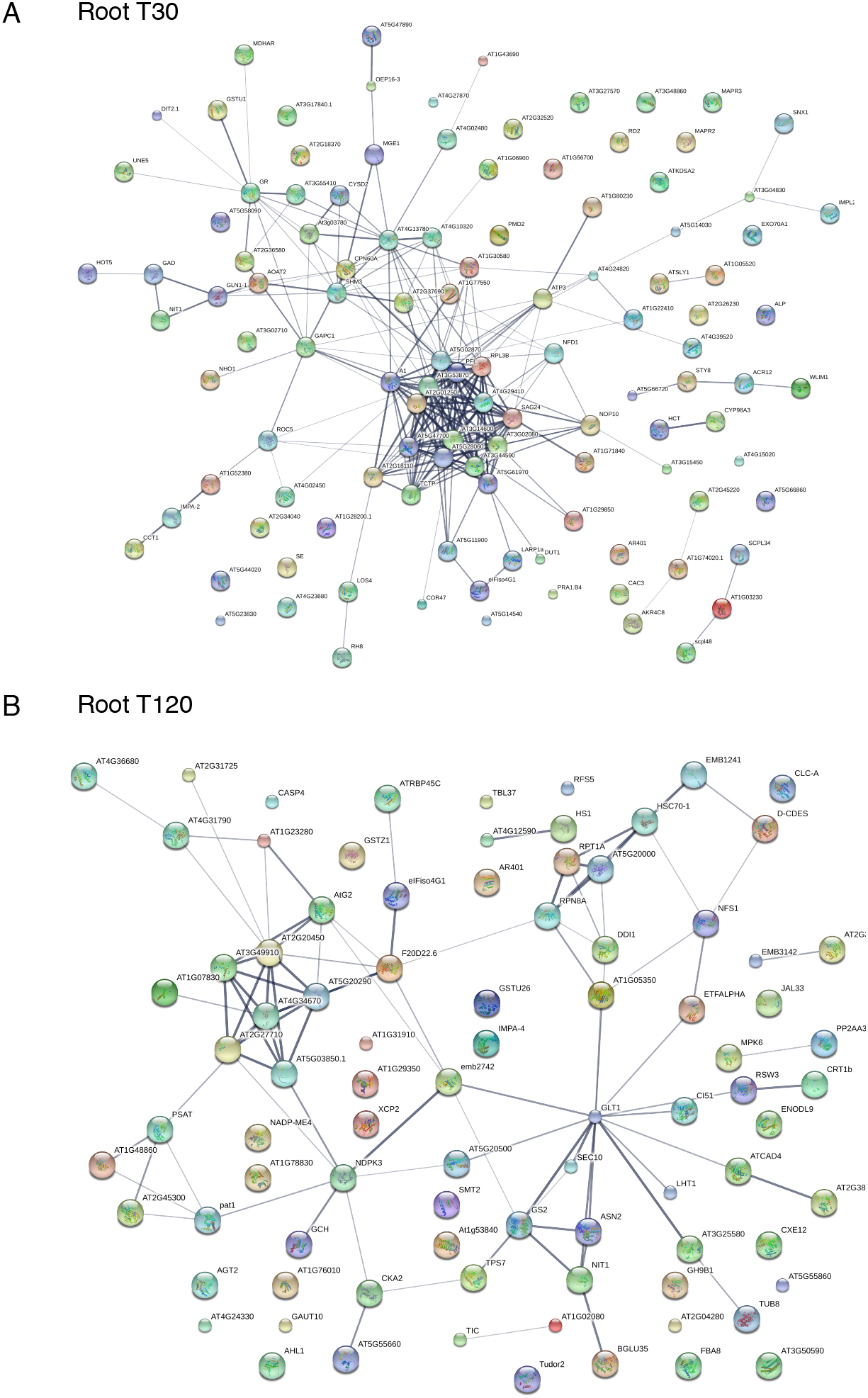
Protein interaction networks generated with root specific auxin regulated proteins are highly connected and enriched for processes such as translation. (A) STRING network of differentially expressed proteins in roots at T30 time point are highly connected with a central hub overrepresented by the term ‘translation’. (A) STRING network of differentially expressed proteins in roots at T120 time point are highly connected but differ in architecture compared to the T30 network.

## Discussion

The effects of auxin on gene regulation have been well appreciated at the transcriptional level. In this study, we describe rapid and quantitative auxin-mediated proteome changes that occur in Arabidopsis seedlings, roots and hypocotyls using quantitative proteomics. These datasets show that auxin regulated proteins belong to diverse functional categories such as abiotic stress, amino acid metabolism, RNA and protein regulation and lipid metabolism. Additionally, they exhibit tissue and temporal specificity. This is consistent with the long-standing notion that auxin drives different developmental outcomes in an organ specific context, thus we propose that these morphogenesis events are shaped by distinct cellular proteomes.

Comparisons between auxin regulated transcriptomes with proteomes showed a striking lack of overlap, suggesting that these levels of gene regulation are not well integrated. In both the root and hypocotyl at T120 min there are 5 proteins in common, including MAP KINASE 6 (MAPK6; At2g43790), which is known to be auxin up-regulated and play roles in root cell division (Contreras-Cornejo et al., 2015; Smékalová et al., 2014).

TRYPTOPHAN BIOSYNTHESIS 1 (TRP1; At5g17990) is also observed in this overlap, which could suggest a feed-forward loop involving up-regulation of auxin biosynthesis following auxin treatment. One of the very few genes that is differentially expressed at both levels in the hypocotyl is PECTIN METHYLESTERASE1 (PME1), which has not directly been linked to auxin signaling per se, but auxin mediated phyllotaxis involves changes in pectin composition in cell walls (Braybrook and Peaucelle, 2013; Peaucelle et al., 2011). Overall, these discordant relationships may result from indirect effects of auxin on various gene regulation events such as mRNA stability, translational control and/or protein degradation. While many of the differentially regulated proteins can be grouped into protein and RNA MapMan bins, there is no clear pattern among up- and downregulated proteins that could satisfactorily explain this observation. Additionally, 5-8% of the auxin regulated proteins reported here are encoded by an upstream open reading frame (uORF) containing gene (Hu et al., 2016) (Table S1). Future studies aimed at examining auxin mediated translational control and protein stability may provide further clarity in this area.

Previous proteome studies based on auxin responses in seedlings and roots involved older seedlings and later time points compared to this study (Slade et al., 2017; Xing and Xue, 2012). Slade et. al., Proteomes 2017 looked at auxin mediated proteome changes in young seedlings 8, 12 and 24 h after treatment; Xing and Xue examined proteome composition in seedlings 6, 12 and 24 h following auxin treatment. Because these are later time points than what we sampled here it is difficult to directly compare the results between these studies, however we did look for proteins in common between these studies and several proteins in common in total that are differentially regulated in the root following auxin treatment (Table S3). Altogether these proteins may represent a set of auxin biomarkers that are rapidly and stably expressed following auxin treatment and are reproducibly detected via peptide mass spectrometry. They include proteins such as SORTING NEXIN 1, CULLIN 3 and NITRILASE; collectively these proteins play important functional roles in various aspects of auxin transport, signaling and biosynthesis (Ambrose et al., 2013; Gusmaroli et al., 2007; Hanzawa et al., 2013; Jaillais et al., 2006; Lehmann et al., 2017; Roberts et al., 2011; Zhang et al., 2013).

Metabolic changes associated with auxin signaling are not well understood and cannot be accurately predicted from transcriptomic data. Quantitative measurements in protein abundance are critical for elucidating what metabolic changes are driven by auxin (Figure 4). Interactions between redox and auxin have recently been considered to be critical for meristem activity (Tognetti et al., 2012, 2017). However, proteins associated with redox were differentially regulated in hypocotyls, roots and seedlings in response to auxin suggesting that alteration of redox state may be a common rapid signaling mechanism that follows early auxin response irrespective of spatial context. Amino acid metabolism was a common category observed among all the datasets (Figure 4), which is consistent with several recently described links between Target of rapamycin (TOR) and auxin (Cai et al., 2017; Deng et al., 2016; Li et al., 2017; Pu et al., 2017; Salem et al., 2017). Proteins associated with lipid metabolism were differentially regulated in hypocotyls, roots and seedlings at all time points, including well characterized acyl-Co-A oxidase (ACX) enzymes that play roles in auxin mediated growth (Adham et al., 2005; Agarwal et al., 2001), suggesting that changes in lipid composition may underlie auxin driven growth processes.

This study sought to provide an extensive description of how auxin signaling influences cellular proteomes in organ specific context. While hundreds of proteins change rapidly in response to auxin in seedlings, we also observed distinct changes within hypocotyl and root tissue. These datasets provide a rich resource for mining novel protein function. In particular, numerous proteins show significant altered abundance levels in a spatiotemporal fashion which makes these excellent candidates for future functional studies. Additionally, these datasets can inform new hypotheses of what biological processes may govern rapid auxin responses downstream of perception, including complex levels of gene regulation and rapid alteration of metabolic states.

## AUTHOR CONTRIBUTIONS

D.R.K, E.J.C., Z.S., S.P.B., and M.E. designed the research. D.R.K, E.J.C., and Z.S. performed the research. D.R.K, J.W.W. and Z.S. analyzed data. D.R.K. wrote the article with input from the other authors.

## Acknowledgments

The authors would like to thank Gloria Muday for sharing microarray data from auxin treated roots associated with Lewis et. al., 2013, Brian Dilkes and Katayoon (Katie) Dehesh for ideas and discussion.

## Supplementary Material

**Table S1.** Auxin responsive proteomes for seedlings, hypocotyls and roots at various time points in Arabidopsis. iTRAQ intensities, log2 ratios and p-values for all proteins detected. Proteins with a p-value ≤ 0.05 were considered to be significant and are listed as separate sheets within the file; those with uORF containing genes are indicated.

**Table S2.** GO enrichment analyses performed on differentially expressed proteins.

**Table S3.** Proteins in common between this study and other published auxin proteome studies.

## References

Adham, A. R., Zolman, B. K., Millius, A., and Bartel, B. (2005). Mutations in Arabidopsis acyl-CoA oxidase genes reveal distinct and overlapping roles in β-oxidation. Plant J. 41, 859–874. doi:10.1111/j.1365-313X.2005.02343.x.

Agarwal, A. K., Qi, Y., Bhat, D. G., Woerner, B. M., and Brown, S. M. (2001). Gene isolation and characterization of two acyl CoA oxidases from soybean with broad substrate specificities and enhanced expression in the growing seedling axis. Plant Mol. Biol. 47, 519–531. doi:10.1023/A:1011825114301.

Ambrose, C., Ruan, Y., Gardiner, J., Tamblyn, L. M., Catching, A., Kirik, V., et al. (2013). CLASP Interacts with Sorting Nexin 1 to Link Microtubules and Auxin Transport via PIN2 Recycling in Arabidopsis thaliana. Dev. Cell 24, 649–659. doi:10.1016/j.devcel.2013.02.007.

Baerenfaller, K., Grossmann, J., Grobei, M. A., Hull, R., Hirsch-Hoffmann, M., Yalovsky, S., et al. (2008). Genome-Scale Proteomics Reveals Arabidopsis thaliana Gene Models and Proteome Dynamics. Science (80-.). 320, 938–941. doi:10.1126/science.1157956.

Bargmann, B. O. R., Vanneste, S., Krouk, G., Nawy, T., Efroni, I., Shani, E., et al. (2014). A map of cell type-specific auxin responses. Mol. Syst. Biol. 9, 688–688. doi:10.1038/msb.2013.40.

Braybrook, S. A., and Peaucelle, A. (2013). Mechano-Chemical Aspects of Organ Formation in Arabidopsis thaliana: The Relationship between Auxin and Pectin. PLoS One 8. doi:10.1371/journal.pone.0057813.

Cai, W., Li, X., Liu, Y., Wang, Y., Zhou, Y., Xu, T., et al. (2017). COP1 Integrates Light Signals to ROP2 for Cell Cycle Activation. Plant Signal. Behav., 00–00. doi:10.1080/15592324.2017.1363946.

Cai, Y., Zhuang, X., Gao, C., Wang, X., and Jiang, L. (2014). The Arabidopsis Endosomal Sorting Complex Required for Transport III Regulates Internal Vesicle Formation of the Prevacuolar Compartment and Is Required for Plant Development. PLANT Physiol. 165, 1328–1343. doi:10.1104/pp.114.238378.

Chapman, E. J., Greenham, K., Castillejo, C., Sartor, R., Bialy, A., Sun, T. ping, et al. (2012). Hypocotyl transcriptome reveals auxin regulation of growth-promoting genes through GA-dependent and -independent pathways. PLoS One 7. doi:10.1371/journal.pone.0036210.

Contreras-Cornejo, H. A., Lopez-Bucio, J. S., Mendez-Bravo, A., Macias-Rodriguez, L., Ramos-Vega, M., Guevara-Garcia, A. A., et al. (2015). Mitogen-Activated Protein Kinase 6 and Ethylene and Auxin Signaling Pathways Are Involved in Arabidopsis Root-System Architecture Alterations by Trichoderma atroviride. Mol. Plant. Microbe. Interact. 28, MPMI01150005R. doi:10.1094/MPMI-01-15-0005-R.

Dai, M., Zhang, C., Kania, U., Chen, F., Xue, Q., Mccray, T., et al. (2012). A PP6-Type Phosphatase Holoenzyme Directly Regulates PIN Phosphorylation and Auxin Efflux in Arabidopsis. Plant Cell 24, 2497–2514. doi:10.1105/tpc.112.098905.

del Pozo, J. C., Dharmasiri, S., Hellmann, H., Walker, L., Gray, W. M., and Estelle, M. (2002). AXR1-ECR1-dependent conjugation of RUB1 to the Arabidopsis Cullin AtCUL1 is required for auxin response. Plant Cell 14, 421–433. doi:10.1105/tpc.010282.

Deng, K., Yu, L., Zheng, X., Zhang, K., Wang, W., Dong, P., et al. (2016). Target of Rapamycin Is a Key Player for Auxin Signaling Transduction in Arabidopsis. Front. Plant Sci. 7. doi:10.3389/fpls.2016.00291.

Dharmasiri, S., Dharmasiri, N., Hellmann, H., and Estelle, M. (2003). The RUB/Nedd8 conjugation pathway is required for early development in Arabidopsis. EMBO J. 22, 1762–1770. doi:10.1093/emboj/cdg190.

Eden, E., Navon, R., Steinfeld, I., Lipson, D., and Yakhini, Z. (2009). GOrilla: a tool for discovery and visualization of enriched GO terms in ranked gene lists. BMC Bioinformatics 10, 48. doi:10.1186/1471-2105-10-48.

Esteve-Bruna, D., Pérez-Pérez, J. M., Ponce, M. R., and Micol, J. L. (2013). incurvata13, a novel allele of AUXIN RESISTANT6, reveals a specific role for auxin and the SCF complex in Arabidopsis embryogenesis, vascular specification, and leaf flatness. Plant Physiol. 161, 1303–20. doi:10.1104/pp.112.207779.

Finet, C., and Jaillais, Y. (2012). AUXOLOGY: When auxin meets plant evo-devo. Dev. Biol. 369, 19–31. doi:10.1016/j.ydbio.2012.05.039.

Franceschini, A., Szklarczyk, D., Frankild, S., Kuhn, M., Simonovic, M., Roth, A., et al. (2013). STRING v9.1: Protein-protein interaction networks, with increased coverage and integration. Nucleic Acids Res. 41. doi:10.1093/nar/gks1094.

Friml, J., Wiśniewska, J., Benková, E., Mendgen, K., and Palme, K. (2002). Lateral relocation of auxin efflux regulator PIN3 mediates tropism in Arabidopsis. Nature 415, 806–809. doi:10.1038/415806a.

Ghaemmaghami, S., Ghaemmaghami, S., Huh, W.-K., Huh, W.-K., Bower, K., Howson, R., et al. (2003). Global analysis of protein expression in yeast. Nature 425, 737–41. doi:10.1038/nature02046.

Ghazalpour, A., Bennett, B., Petyuk, V. A., Orozco, L., Hagopian, R., Mungrue, I. N., et al. (2011). Comparative analysis of proteome and transcriptome variation in mouse. PLoS Genet. 7. doi:10.1371/journal.pgen.1001393.

Gonzalez-Lamothe, R., El Oirdi, M., Brisson, N., and Bouarab, K. (2012). The Conjugated Auxin Indole-3-Acetic Acid-Aspartic Acid Promotes Plant Disease Development. PLANT CELL ONLINE 24, 762–777. doi:10.1105/tpc.111.095190.

Gusmaroli, G., Figueroa, P., Serino, G., and Deng, X. W. (2007). Role of the MPN subunits in COP9 signalosome assembly and activity, and their regulatory interaction with Arabidopsis Cullin3-based E3 ligases. Plant Cell 19, 564–81. doi:10.1105/tpc.106.047571.

Hanzawa, T., Shibasaki, K., Numata, T., Kawamura, Y., Gaude, T., and Rahman, A. (2013). Cellular Auxin Homeostasis under High Temperature Is Regulated through a SORTING NEXIN1-Dependent Endosomal Trafficking Pathway. Plant Cell 25, 3424–3433. doi:10.1105/tpc.113.115881.

Harper, R. M., Stowe-Evans, E. L., Luesse, D. R., Muto, H., Tatematsu, K., Watahiki, M. K., et al. (2000). The NPH4 locus encodes the auxin response factor ARF7, a conditional regulator of differential growth in aerial Arabidopsis tissue. Plant Cell 12, 757–70. doi:10.1105/tpc.12.5.757.

Horiguchi, G., Van Lijsebettens, M., Candela, H., Micol, J. L., and Tsukaya, H. (2012). Ribosomes and translation in plant developmental control. Plant Sci. 191–192, 24. 34. doi:10.1016/j.plantsci.2012.04.008.

Hu, Q., Merchante, C., Stepanova, A., Alonso, J., and Heber, S. (2016). Genome-wide Search for Translated Upstream Open Reading Frames in Arabidopsis thaliana. IEEE Trans. Nanobioscience 15, 148–157. doi:10.1109/TNB.2016.2516950.

Jaillais, Y., Fobis-Loisy, I., Miège, C., Rollin, C., and Gaude, T. (2006). AtSNX1 defines an endosome for auxin-carrier trafficking in Arabidopsis. Nature 443, 106–109. doi:10.1038/nature05046.

Laskowski, M., Biller, S., Stanley, K., Kajstura, T., and Prusty, R. (2006). Expression profiling of auxin-treated Arabidopsis roots: Toward a molecular analysis of lateral root emergence. Plant Cell Physiol. 47, 788–792. doi:10.1093/pcp/pcj043.

Lehmann, T., Janowitz, T., Sánchez-Parra, B., Alonso, M.-M. P., Trompetter, I., Piotrowski, M., et al. (2017). Arabidopsis NITRILASE 1 Contributes to the Regulation of Root Growth and Development through Modulation of Auxin Biosynthesis in Seedlings. Front. Plant Sci. 8. doi:10.3389/fpls.2017.00036.

Lewis, D. R., Olex, A. L., Lundy, S. R., Turkett, W. H., Fetrow, J. S., and Muday, G. K. (2013). A Kinetic Analysis of the Auxin Transcriptome Reveals Cell Wall Remodeling Proteins That Modulate Lateral Root Development in Arabidopsis. Plant Cell 25, 3329–3346. doi:10.1105/tpc.113.114868.

Li, X., Cai, W., Liu, Y., Li, H., Fu, L., Liu, Z., et al. (2017). Differential TOR activation and cell proliferation in Arabidopsis root and shoot apexes. Proc. Natl. Acad. Sci. 114, 2765–2770. doi:10.1073/pnas.1618782114.

Mattei, B., Sabatini, S., and Schininà, M. E. (2013). Proteomics in deciphering the auxin commitment in the Arabidopsis thaliana root growth. J. Proteome Res. 12, 4685–4701. doi:10.1021/pr400697s.

Nemhauser, J. L., Hong, F., and Chory, J. (2006). Different Plant Hormones Regulate Similar Processes through Largely Nonoverlapping Transcriptional Responses. Cell 126, 467–475. doi:10.1016/j.cell.2006.05.050.

Nishimura, T., Wada, T., Yamamoto, K. T., and Okada, K. (2005). The Arabidopsis STV1 protein, responsible for translation reinitiation, is required for auxin-mediated gynoecium patterning. Plant Cell 17, 2940–2953. doi:10.1105/tpc.105.036533.

Overvoorde, P. J., Okushima, Y., Alonso, J. M., Chan, A., Chang, C., Ecker, J. R., et al. (2005). Functional genomic analysis of the AUXIN/INDOLE-3-ACETIC ACID gene family members in Arabidopsis thaliana. Plant Cell 17, 3282–3300. doi:10.1105/tpc.105.036723.

Paponov, I. A., Teale, W. D., Trebar, M., Blilou, I., and Palme, K. (2005). The PIN auxin efflux facilitators: Evolutionary and functional perspectives. Trends Plant Sci. 10, 170–177. doi:10.1016/j.tplants.2005.02.009.

Peaucelle, A., Braybrook, S. A., Le Guillou, L., Bron, E., Kuhlemeier, C., and Höfte, H. (2011). Pectin-induced changes in cell wall mechanics underlie organ initiation in Arabidopsis. Curr. Biol. 21, 1720–1726. doi:10.1016/j.cub.2011.08.057.

Ponnala, L., Wang, Y., Sun, Q., and Van Wijk, K. J. (2014). Correlation of mRNA and protein abundance in the developing maize leaf. Plant J. 78, 424–440. doi:10.1111/tpj.12482.

Pozo, J. C. (1998). The Ubiquitin-Related Protein RUB1 and Auxin Response in Arabidopsis. Science (80-.). 280, 1760–1763. doi:10.1126/science.280.5370.1760.

Provart, N. J., Gil, P., Chen, W., Han, B., Chang, H.-S., Wang, X., et al. (2003). Gene expression phenotypes of Arabidopsis associated with sensitivity to low temperatures. Plant Physiol. 132, 893–906. doi:10.1104/pp.103.021261.

Pu, Y., Luo, X., and Bassham, D. C. (2017). TOR-Dependent and -Independent Pathways Regulate Autophagy in Arabidopsis thaliana. Front. Plant Sci. 8. doi:10.3389/fpls.2017.01204.

Roberts, D., Pedmale, U. V., Morrow, J., Sachdev, S., Lechner, E., Tang, X., et al. (2011). Modulation of Phototropic Responsiveness in Arabidopsis through Ubiquitination of Phototropin 1 by the CUL3-Ring E3 Ubiquitin Ligase CRL3NPH3. Plant Cell 23, 3627–3640. doi:10.1105/tpc.111.087999.

Rosado, A., Li, R., van de Ven, W., Hsu, E., and Raikhel, N. V (2012). Arabidopsis ribosomal proteins control developmental programs through translational regulation of auxin response factors. Proc. Natl. Acad. Sci. U. S. A. 109, 19537–44. doi:10.1073/pnas.1214774109.

Salem, M. A., Li, Y., Wiszniewski, A., and Giavalisco, P. (2017). Regulatory-Associated Protein of TOR (RAPTOR) alters the hormonal and metabolic composition of Arabidopsis seeds controlling seed morphology, viability and germination potential. Plant J. doi:10.1111/tpj.13667.

Schwanhausser, B., Busse, D., Li, N., Dittmar, G., Schuchhardt, J., Wolf, J., et al. (2011). Global quantification of mammalian gene expression control. Nature 473, 337–342. doi:10.1038/nature10098.

Slade, W., Ray, W., Hildreth, S., Winkel, B., and Helm, R. (2017). Exogenous Auxin Elicits Changes in the Arabidopsis thaliana Root Proteome in a Time-Dependent Manner. Proteomes 5, 16. doi:10.3390/proteomes5030016.

Smékalová, V., Luptovčiak, I., Komis, G., Šamajová, O., Ovečka, M., Doskočilová, A., et al. (2014). Involvement of YODA and mitogen activated protein kinase 6 in Arabidopsis post-embryogenic root development through auxin up-regulation and cell division plane orientation. New Phytol. 203, 1175–1193. doi:10.1111/nph.12880.

Staswick, P. E. (2005). Characterization of an Arabidopsis Enzyme Family That Conjugates Amino Acids to Indole-3-Acetic Acid. PLANT CELL ONLINE 17, 616–627. doi:10.1105/tpc.104.026690.

Stepanova, A. N., Yun, J., Likhacheva, A. V., and Alonso, J. M. (2007). Multilevel Interactions between Ethylene and Auxin in Arabidopsis Roots. PLANT CELL ONLINE 19, 2169–2185. doi:10.1105/tpc.107.052068.

Strader, L. C., and Zhao, Y. (2016). Auxin perception and downstream events. Curr. Opin. Plant Biol. 33, 8–14. doi:10.1016/j.pbi.2016.04.004.

Swarup, R., Kargul, J., Marchant, A., Zadik, D., Rahman, A., Mills, R., et al. (2004). Structure-Function Analysis of the Presumptive Arabidopsis Auxin Permease AUX1. Plant Cell 16, 3069–3083. doi:10.1105/tpc.104.024737.

Szklarczyk, D., Franceschini, A., Wyder, S., Forslund, K., Heller, D., Huerta-Cepas, J., et al. (2015). STRING v10: Protein-protein interaction networks, integrated over the tree of life. Nucleic Acids Res. 43, D447–D452. doi:10.1093/nar/gku1003.

Szklarczyk, D., Morris, J. H., Cook, H., Kuhn, M., Wyder, S., Simonovic, M., et al. (2017). The STRING database in 2017: quality-controlled protein-protein association networks, made broadly accessible. Nucleic Acids Res. 45, D362–D368. doi:10.1093/nar/gkw937.

Tian, C. E., Muto, H., Higuchi, K., Matamura, T., Tatematsu, K., Koshiba, T., et al. (2004). Disruption and overexpression of auxin response factor 8 gene of Arabidopsis affect hypocotyl elongation and root growth habit, indicating its possible involvement in auxin homeostasis in light condition. Plant J. 40, 333–343. doi:10.1111/j.1365-313X.2004.02220.x.

Tognetti, V. B., Bielach, A., and Hrtyan, M. (2017). Redox regulation at the site of primary growth: Auxin, cytokinin and ROS crosstalk. Plant. Cell Environ., 1–20. doi:10.1111/pce.13021.

Tognetti, V. B., Mühlenbock, P., and van Breusegem, F. (2012). Stress homeostasis - the redox and auxin perspective. Plant, Cell Environ. 35, 321–333. doi:10.1111/j.1365-3040.2011.02324.x.

Ugartechea-Chirino, Y., Swarup, R., Swarup, K., Péret, B., Whitworth, M., Bennett, M., et al. (2010). The AUX1 LAX family of auxin influx carriers is required for the establishment of embryonic root cell organization in Arabidopsis thaliana. Ann. Bot. 105, 277–289. doi:10.1093/aob/mcp287.

Vogel, C., de Sousa Abreu, R., Ko, D., Le, S.-Y., Shapiro, B. A., Burns, S. C., et al. (2010). Sequence signatures and mRNA concentration can explain two-thirds of protein abundance variation in a human cell line. Mol. Syst. Biol. 6. doi:10.1038/msb.2010.59.

Waese, J., and Provart, N. J. (2016). The Bio-Analytic Resource: Data visualization and analytic tools for multiple levels of plant biology. Curr. Plant Biol. 7–8, 2–5. doi:10.1016/j.cpb.2016.12.001.

Walley, J. W., Sartor, R. C., Shen, Z., Schmitz, R. J., Wu, K. J., Urich, M. A., et al. (2016). Integration of omic networks in a developmental atlas of maize. Science 353, 814–8. doi:10.1126/science.aag1125.

Walley, J. W., Shen, Z., Sartor, R., Wu, K. J., Osborn, J., Smith, L. G., et al. (2013). Reconstruction of protein networks from an atlas of maize seed proteotypes. Proc. Natl. Acad. Sci. 110, E4808–E4817. doi:10.1073/pnas.1319113110.

Walley, J., Xiao, Y., Wang, J.-Z., Baidoo, E. E., Keasling, J. D., Shen, Z., et al. (2015). Plastid-produced interorgannellar stress signal MEcPP potentiates induction of the unfolded protein response in endoplasmic reticulum. Proc. Natl. Acad. Sci. 112, 6212–6217. doi:10.1073/pnas.1504828112.

Washburn, M. P., Koller, A., Oshiro, G., Ulaszek, R. R., Plouffe, D., Deciu, C., et al. (2003). Protein pathway and complex clustering of correlated mRNA and protein expression analyses in Saccharomyces cerevisiae. Proc. Natl. Acad. Sci. 100, 3107–3112. doi:10.1073/pnas.0634629100.

Watahiki, M. K., Tatematsu, K., Fujihira, K., Yamamoto, M., and Yamamoto, K. T. (1999). The MSG1 and AXR1 genes of Arabidopsis are likely to act independently in growth-curvature responses of hypocotyls. Planta 207, 362–369. doi:10.1007/s004250050493.

Weijers, D., and Wagner, D. (2016). Transcriptional Responses to the Auxin Hormone. Annu. Rev. Plant Biol. 67, 539–574. doi:10.1146/annurev-arplant-043015-112122.

Withers, J. C., Shipp, M. J., Rupasinghe, S. G., Sukumar, P., Schuler, M. A., Muday, G. K., et al. (2013). Gravity persistent signal 1 (GPS1) reveals novel cytochrome P450s involved in Gravitropism. Am. J. Bot. 100, 183–193. doi:10.3732/ajb.1200436.

Xing, M., and Xue, H. (2012). A proteomics study of auxin effects in Arabidopsis thaliana. Acta Biochim. Biophys. Sin. (Shanghai). 44, 783–796. doi:10.1093/abbs/gms057.

Zhang, H., Zhou, H., Berke, L., Heck, A. J. R., Mohammed, S., Scheres, B., et al. (2013). Quantitative Phosphoproteomics after Auxin-stimulated Lateral Root Induction Identifies an SNX1 Protein Phosphorylation Site Required for Growth. Mol. Cell. Proteomics 12, 1158–1169. doi:10.1074/mcp.M112.021220.

Zhou, F., Roy, B., and von Arnim, A. G. (2010). Translation reinitiation and development are compromised in similar ways by mutations in translation initiation factor eIF3h and the ribosomal protein RPL24. BMC Plant Biol. 10, 193. doi:10.1186/1471-2229-10-193.

